# MarrowDLD: a microfluidic method for label-free retrieval of fragile bone marrow cells

**DOI:** 10.1101/2023.09.22.558939

**Authors:** Gloria Porro, Rita Sarkis, Clara Obergozo, Lucie Godot, Francesco Amato, Magali Humbert, Olaia Naveiras, Carlotta Guiducci

## Abstract

Functional bone marrow studies have focused primarily on hematopoietic progenitors, leaving limited knowledge about other fragile populations, such as bone marrow adipocytes (BMAds) and megakaryocytes. The isolation of these cells is challenging due to rupture susceptibility and large size. We introduce here a label-free cytometry microsystem, MarrowDLD, based on deterministic lateral displacement. MarrowDLD enables the isolation of fragile cells based on intrinsic size properties while preserving their viability and functionality. Bone marrow adipocytes, obtained from mouse and human stromal line differentiation, as well as megakaryocytes, from primary human CD34+ hematopoietic stem and progenitor cells, were used for validation. Precise micrometer-range separation cutoffs were adapted for each cell type. Cells were sorted directly in culture media, without pre-labeling steps, and with real-time imaging for quality control. At least 10^6^ cells were retrieved intact per sorting round. Our method outperformed two FACS instruments in purity and yield, particularly for large cell size fractions. MarrowDLD represents a non-destructive sorting tool for fragile cells, facilitating the separation of phenotypically pure populations of BMAds and megakaryocytes to further investigate their physiological and pathological roles.

## Introduction

The bone marrow (BM) constitutes the primary site of hematopoiesis, where maturing hematopoietic cells and supporting stromal cells coexist within a complex microenvironment ensuring the tightly regulated production of up to 10^12^ blood cells daily [1]. However, certain BM cell types, namely bone marrow adipocytes (BMAds) and megakaryocytes (MK), have eluded comprehensive characterizations due to their fragile nature and considerable size when fully mature, complicating their isolation via conventional methods such as flow cytometry and fluorescence-activated cell sorting (FACS) [2]. Consequently, they are often underrepresented or absent in critical studies such as single-cell RNA sequencing-based atlases [3]–[6].

Mature megakaryocytes are large cells, 50-100 μm in diameter [7], which produce platelets, the cell fragments mediating blood clotting. Despite efforts to decode the impact of diverse MK phenotypes on platelet generation and hematopoietic progenitor fate, the lack of robust functional studies has hindered conclusive determinations [7]. While *in vitro* differentiation of megakaryocytes from hematopoietic stem cells offers insights into megakaryopoiesis and platelet function[8], the isolation of high-purity, viable MK subpopulations for mechanistic investigations remains a hurdle [9]–[11]. The exigency of separating mature MKs from their precursors is critical to define differentiation requirements and optimize *ex vivo* platelet production. Although megakaryocytes express specific surface markers facilitating their separation through flow cytometry [7], [12], [13], their rarity and susceptibility to shear stresses limit their isolation. Indeed, FACS, the gold-standard method for high-throughput cell sorting, loses efficacy when applied to cells with either large size, inherent fragility, and/or high buoyancy [14]–[16].

Sorting bone marrow adipocytes is even more restrictive than MKs. BMAds constitute the most frequent BM stromal cells in larger mammals [17], varying in size from 30-40 μm in mice to 80-100 μm in humans [18]–[20]. Historically regarded as passive space fillers, BMAds are now recognized as regulators of energy storage, bone metabolism, and hematopoiesis [21]– [23]. As the interest in their composition, function, and heterogeneity has increased [24]–[27], their difficult isolation and preservation upon sorting have hindered research progress [2]. Additionally, since fully-lipidated mature adipocytes lack specific surface markers, neutral lipid dyes are often used for their FACS-based sorting. Yet, the labeling steps are time-consuming and affect cell viability. Notably, lipid dyes are not exclusive to mature adipocytes since they also stain progenitor cell membranes and early differentiation stages [14], [28]–[30]. FACS sorting combining lipid staining with forward scatter (FSC) and side scatter (SSC), which reflects cell size and granularity, is a common strategy [15], [31]–[34]. Recently, a size-based FACS protocol relying solely on FSC/SSC parameters has been proposed to isolate large unilocular mature adipocytes, albeit restricted to fixed cells and requiring components often unavailable in standard instruments [14]. Given these limitations, buoyancy-based isolation protocols have emerged as an alternative to FACS for the isolation of adipocytes as heterogeneous populations of different sizes and lipid contents [19]. An alternative strategy to separate live BMAds based on size, and thus also to lipid content, would greatly complement the need for homogeneous populations. This approach could overcome the non-specificity of lipid dyes while simplifying sample preparation and preserving cell viability across all maturation stages.

Microfluidic cell separation techniques offer a promising alternative to FACS sorting. Specifically, continuous-flow microfluidic systems are operated at significantly lower pressures than flow cytometry, minimizing shear stresses. Label-free approaches have been implemented with various microfluidic configurations, such as deterministic lateral displacement (DLD). DLD consists of a flow-through microchamber with arrayed micropillars that physically displace large particles, enabling their separation [35]–[38]. Notably, the distances between DLD micropillars are wider than the particles in the processed sample, avoiding compressing even the largest particles in the mixture. Previous research on DLD sorting successfully showed the separation of red and white blood cells from whole blood, achieving post-sorting viability exceeding 90% [39], [40]. Other DLD studies showed the enrichment of human BM skeletal stem cells from expansion of blood extracts [41] and isolation of circulating tumor cells from undiluted blood [42]–[44], achieving throughputs of up to 10^6^ cells per second [43]. A noteworthy commercial DLD implementation is the Curate® Cell Processing System from CurateBio [45], [46], which sorts white blood cells from plasma at 400 mL/hour by running multiple separation microchambers in parallel.

Here, we introduce a novel DLD architecture, MarrowDLD, specifically designed to sort large bone marrow fragile cells, taking advantage of the characteristic size increase that accompanies cell differentiation of BM lineages, whose fragility has greatly limited comprehensive studies. Concretely, MarrowDLD is a fluid dynamic DLD microsystem purifying mature BMAds and MKs from progenitor cells and early stages of differentiation, exclusively based on their size difference. Through extensive testing on BMAds from mouse and human cell lines, as well as primary MKs, we demonstrate effective separation without compromising viability or functionality, enabling cell culture post-sorting. We validate the phenotype of the isolated fractions and compare our method to FACS in terms of purity and functional cell yield. This microfluidic system represents the first non-destructive, phenotype-based label-free sorting method of pure fractions of mature BM adipocytes and megakaryocytes with a precisely defined size range.

## Results

### High-purity isolation of mature OP9 adipocytes

The bone marrow (**Fig. 1A**) is a complex tissue with diverse cellular species. In addition to hematopoietic cells, it includes MKs and BMAds, both characterized by large size and fragility in suspension. To develop our sorting device, we first utilized OP9 cells, a line of bone marrow-derived mouse stromal cells known as a robust adipogenesis model [47], [48]. After *in vitro* adipocytic-induced differentiation, OP9 progenitors accumulated lipid droplets and underwent the expected size increase (**Fig. 1B**, in suspension). **Fig. 1C** shows pre- and post-differentiation OP9 cultures. The differentiation outcome depended on passage number and confluency at induction, thus induced-OP9 samples resulted in varying ratios of differentiation stages, from progenitors to mature adipocytes, ranging from 7 μm to 40 μm in size.

**Figure 1:**
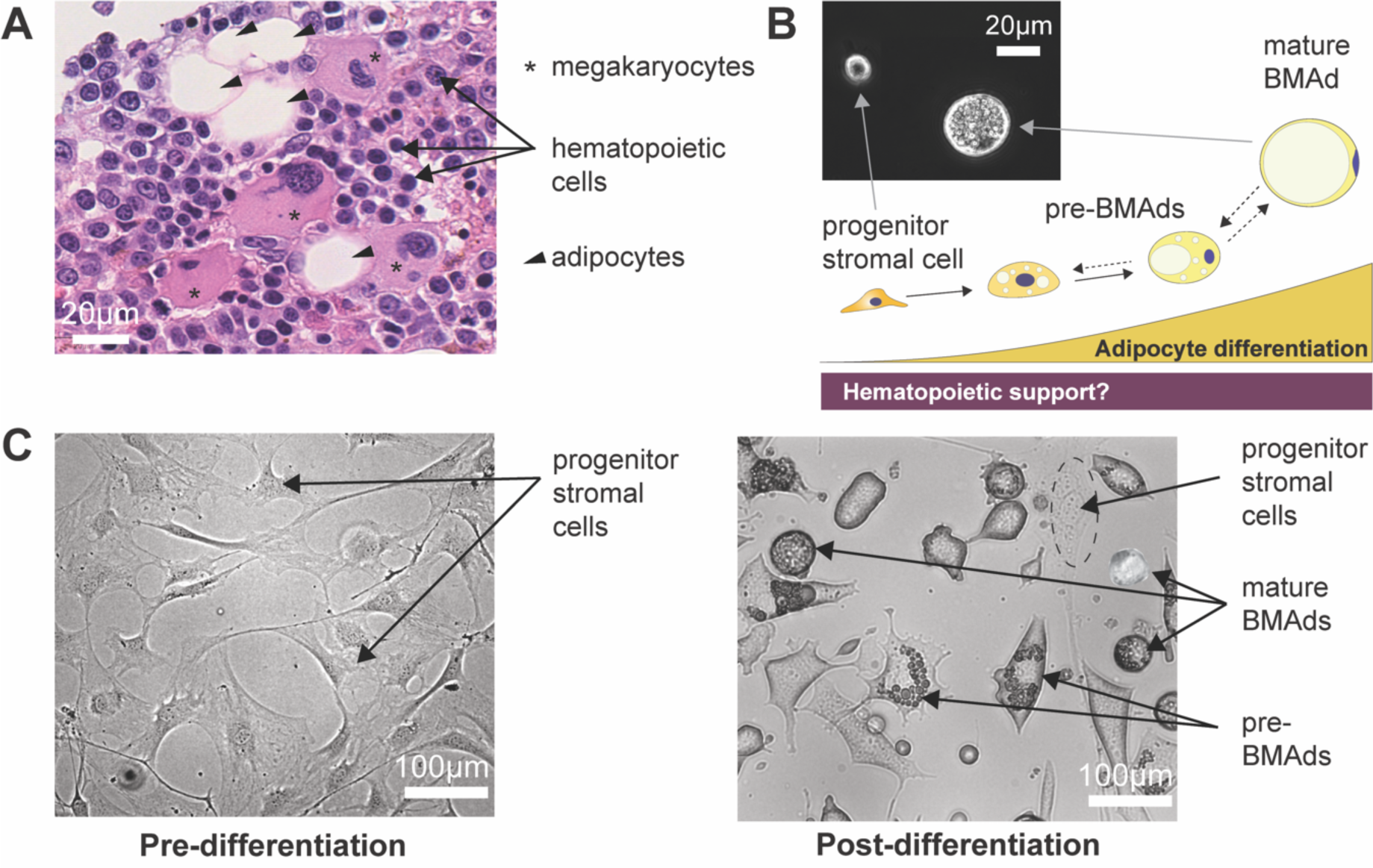
Bone Marrow niche and Heterogeneous Adipocytic Differentiation in vitro. (A) Hematoxylin and eosin-stained slide of human bone marrow (BM) trephine biopsy imaged at 40X magnification using a Nanozoomer S60, revealing hematopoietic cells (labeled with arrows), megakaryocytes (labeled with asterisks), and bone marrow adipocytes (BMAds) (labeled with arrow heads). (B) Schematic representation of the BMAd differentiation axis as a study model for the relationship between hematopoiesis and adipogenesis. Inset: phase contrast imaging of cells in suspension at the progenitor and mature BMAd stages, highlighting the size difference between these two populations. (C) OP9 progenitor cells seeded at 20,000 cells/cm^2^ show a homogeneous fibroblastic-like cell structure in adherent cultures. Following *in vitro-induced* adipocytic differentiation (6 days), the sample contains various stages of maturation: progenitors (fibroblast morphology), pre-BMAds (cells with limited lipid droplet accumulation), and mature BMAds (round cells filled with lipids).

The microchip layout and SEM images of the sorting module are presented in **Fig. 2A-B**, and supplementary videos (**SV1-2**) show the microchip operation. **Fig. 2C-D** depict the behavior of microbeads within MarrowDLD and the sorting workflow for adipocytic cultures. **Figure 3A** shows a sample of OP9-derived adipocytic culture (induced-OP9 [48]) transiting the MarrowDLD sorter at the inlet and outlet. Specifically, a single-cell suspension was continuously injected and focused using lateral sheath flows (**Fig. 3A**, inlet). Stirring the sample prevented bias due to cell buoyancy and an embedded filtering system at the inlet tube reduced doublets and clusters (**Fig. 2D**).

**Figure 2:**
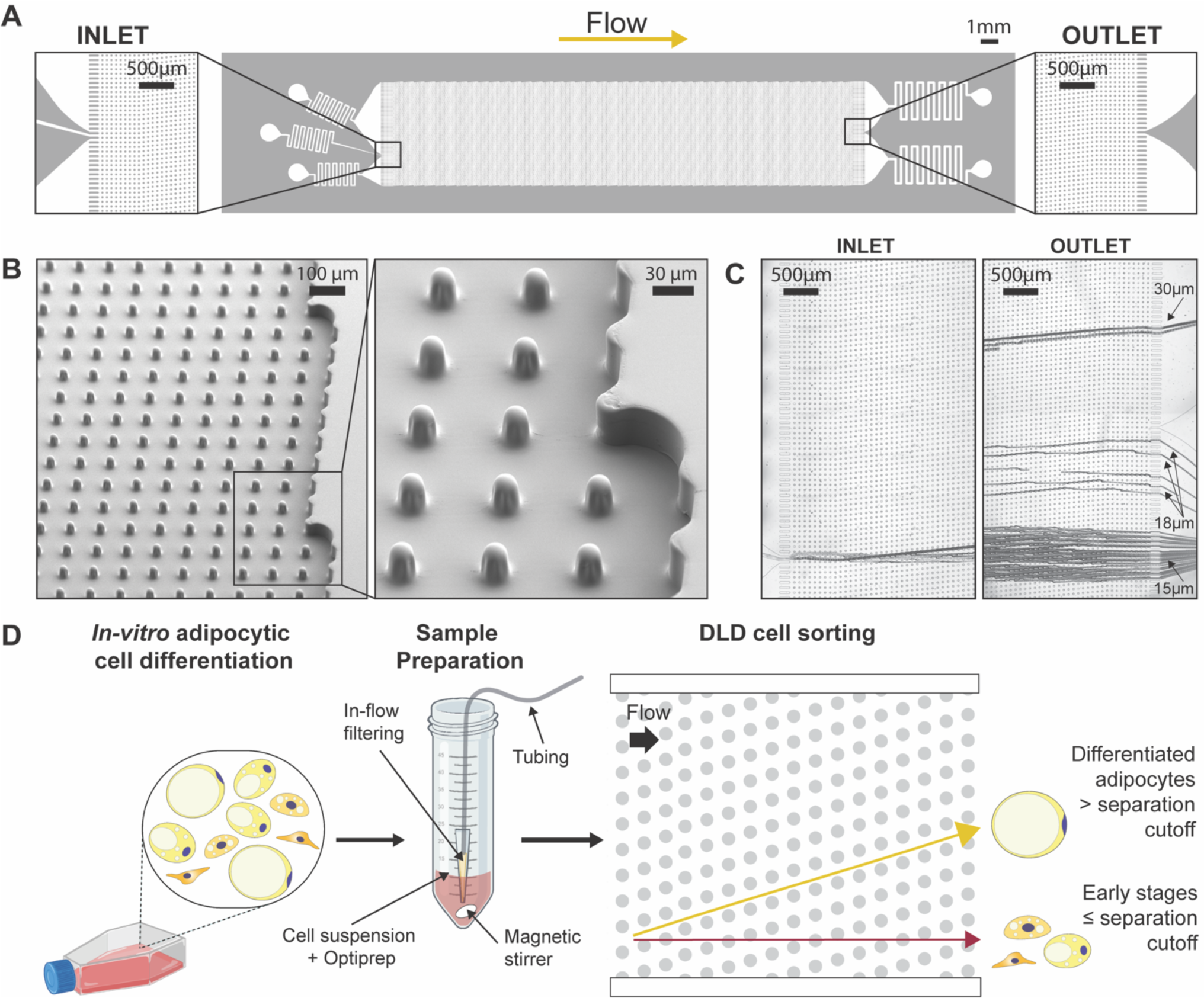
MarrowDLD device and operation. (A) MarrowDLD chip: insights on inlet and outlet regions terminated with 10 rows of straight pillars respectively before and after the MarrowDLD array active region (tilted array). (B) Scanning electron microscopy images of the MarrowDLD array chip with a 19 μm separation cutoff (critical size). (C) Inlet and outlet trajectories of polystyrene microbeads of 15, 18, and 20 μm in size transiting a MarrowDLD chip with 19 μm critical size. (D) Experimental workflow to sort differentiated adipocytes by MarrowDLD. After adipocytic differentiation in vitro, the sample contains a mixture of progenitor cells, early stages of differentiation, and mature adipocytes. After trypsinization, the cellular sample is suspended in its original culture media supplemented with Optiprep, then placed at the inlet reservoir connected to the tubing responsible for injecting the sample into the MarrowDLD device. A custom in-flow filtering system embedded in the tubing ensures the injection of a single-cell suspension, and the sample is continuously stirred to achieve homogeneity. Within the sorting module, mature adipocytes larger than the critical size for separation should be forced to follow the array angle (displacement mode), allowing for their isolation. Conversely, early stages of differentiation and progenitors should move parallel to the flow (zig-zag mode). The two fractions are thus predicted to be physically separated and can be collected at distinct outlets.

**Figure 3:**
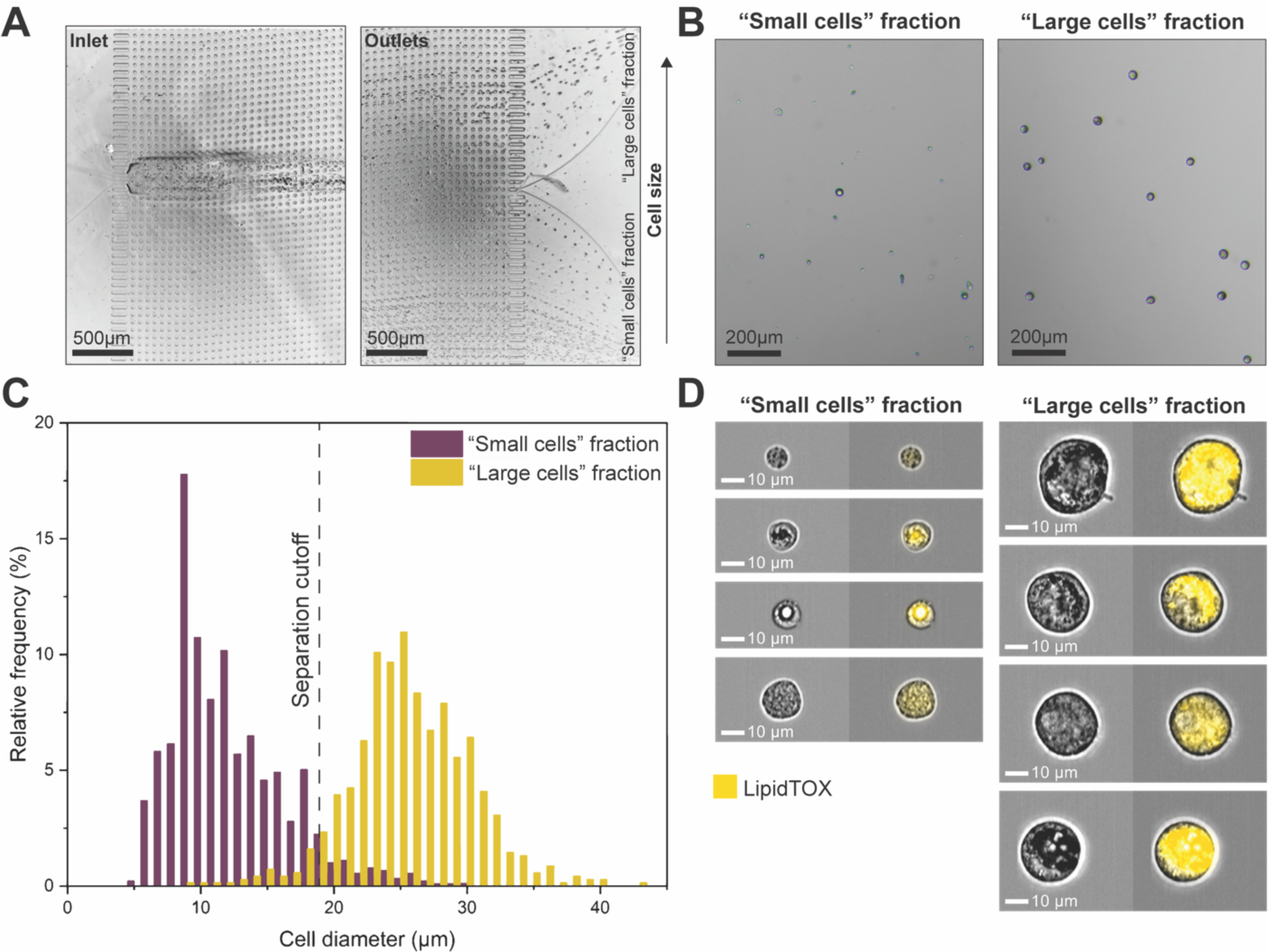
Fluid Dynamic MarrowDLD Sorting of Induced-OP9 Cells. (A) Trajectories of cells entering (inlet) and exiting (outlets) the microfluidic system. The test involved injecting induced-OP9 cells after 6 days of differentiation at a concentration of 500,000 cells/mL. (B) Phase contrast microscopy of outlet fractions after MarrowDLD sorting an induced-OP9 mixture. (C) Cell diameter distributions of outlet fractions (« small cells » fraction, violet; « large cells » fraction, yellow) after sorting induced-OP9s by MarrowDLD with a 19 µm separation cutoff. Phase contrast microscopy of cells collected at the outlets and replated after sorting as in (B) enabled cell size quantification in QuPath0.3.2. Data are displayed as relative frequencies over 150 cells per fraction from n=4 sorting experiments. (D) ImageStream flow cytometry imaging (brightfield and LipidTOX-stained) of cells from the two fractions separated by MarrowDLD.

The MarrowDLD sorting array is designed to retrieve two fractions based on a separation cutoff size determined by the array periodicity and gap between pillars. The injected induced-OP9 single-cell suspension comprised a continuous distribution of cell sizes. As expected from the design, cells exited the array spreading across the width of the main channel, which subdivides downstream into two subchannels denominated « small cells » and « large cells » fractions (**Fig. 3A**, outlets). Notably, the « small cells » outlet collected more events per unit time, including small debris and contaminations, whereas the « large cells » outlet was highly purified (**Fig. 3B**).

We designed and tested MarrowDLD sorting arrays with different separation cutoffs (DLD critical sizes). We fabricated PDMS replicas of the microfluidic devices and measured the pillar gap by surface profilometry. The nominal separation cutoff was calculated based on the general DLD model [39]. We first characterized the sorting modules by processing polystyrene microbeads of different sizes (**Fig. 2C**). Subsequently, we tested several array geometries and observed the sorted fractions by phase contrast microscopy, finding that a 19 μm separation cutoff was ideal for isolating mature OP9 BMAds (**Fig. 3C**). The gap of 42 ± 1 μm in this array was sufficient to preserve intact mature adipocytes with minimal blockage (few units per sorting run).

Specifically, **Figure 3C** shows size distributions of the two output fractions after sorting induced-OP9s by MarrowDLD with a 19 µm separation cutoff. Cells collected at the outlet reservoirs were imaged by phase contrast microscopy, and QuPath0.3.2 software^43^ was used for cell diameter quantification (n=4 independent experiments with 150 cells per fraction). We found that 97 ± 2% of cells within the « large cells » fractions were above the 19 μm cutoff, representing an approximately three-fold enrichment compared to the original OP9-induced adipocytic sample (with only 38% above 19 μm). Consistently, the « small cells » fractions contained 96% ± 3% of cells below the separation cutoff. Independent sorting experiments through different MarrowDLD devices showed reproducible purity for both fractions, independent from the original mixture composition. The combined size distribution of the sorted fractions matches that of the original sample, indicating the preservation of cell size dynamics. **Figure S1** shows the sorting outcome for a device with a separation cutoff of 24 μm, achieving 90-93% purity over the « large cells » fraction for two independent sorting experiments. It is worth noting that we could reliably retrieve intact BMAds above 35 μm in diameter, the size of the largest adipocytes found in the murine BM, as defined within the intact tissue.

To validate our findings, we observed the populations sorted by MarrowDLD by ImageStream flow cytometry imaging (**Fig. 3D**). LipidTOX, a lipophilic stain commonly used for adipocyte FACS sorting, was used to tag the lipid-laden cells. Indeed, all cells within the « large cells » fraction exhibited positive LipidTOX staining, indicating significant lipid accumulation typical of mature BMAds. Large adipocytes and diverse levels of lipid drop coalescence were observed. Interestingly, some cells in the « small cells » fraction also displayed LipidTOX positivity at varying intensities. Some pre-BMAds showed small lipid droplets coalescing, producing a significant LipidTOX signal even if not fully differentiated. These results suggest that BMAds sorting based solely on lipid staining intensity may not be sufficient to differentiate between adipocyte maturation stages. Gently purifying adipocyte populations with a precisely defined size range can help to overcome this limitation.

### Phenotype of the different fractions after sorting induced-OP9 adipocytes

In addition to ImageStream flow cytometry imaging, BODIPY staining and phase contrast microscopy were used to interrogate the phenotype and neutral lipid content of the cells in the small and large cell fractions as compared to the unsorted mix (**Fig. S2A**). To confirm the presence of mature BMAds in the « large cells » fraction and morphologically discern BMAds from their progenitors or intermediately differentiated cells, we employed BODIPY fluorescence stain, indicative of cellular lipidic content and thus of the extent of differentiation. We found the great majority of cells in the « large cells » sorted fraction to stain for BODIPY (**Fig. S2C-D**). Conversely, a very small fraction of cells within the « small cells » fraction was stained for BODIPY, confirming the enrichment of smaller, non-lipidated cells (**Fig. S2B**). Overall, we thus concluded that MarrowDLD sorting of induced-OP9 adipocytic cell suspensions could successfully separate large, lipidated, intact BMAd at high purity from unlipidated or adipogenesis-refractory precursors as determined by the phenotype and lipid content of the sorted cells.

### Viability and functionality of sorted induced-OP9 adipocytes

Next, we evaluated the viability of induced-OP9s before and after MarrowDLD sorting (n=15, with approx. 100 cells per experiment) (**Fig. 4A**). Cell viability was measured using Trypan Blue and a hemocytometer. Sample filtering and Optiprep addition did not significantly influence cell viability (pre-sorting viability: 91%±7%; post-sample preparation viability: 91%±8%). Cells remaining in the input vial after the sorting process exhibited comparable viability (unsorted leftover viability: 86%±8%). As for the sorted cells, a significant but modest decrease in viability (82%±6%) was observed as compared to the pre-sorting sample, but the difference was no longer significant when compared to the unsorted leftover. Overall, viability never fell below 70% for all sorting experiments performed.

**Figure 4:**
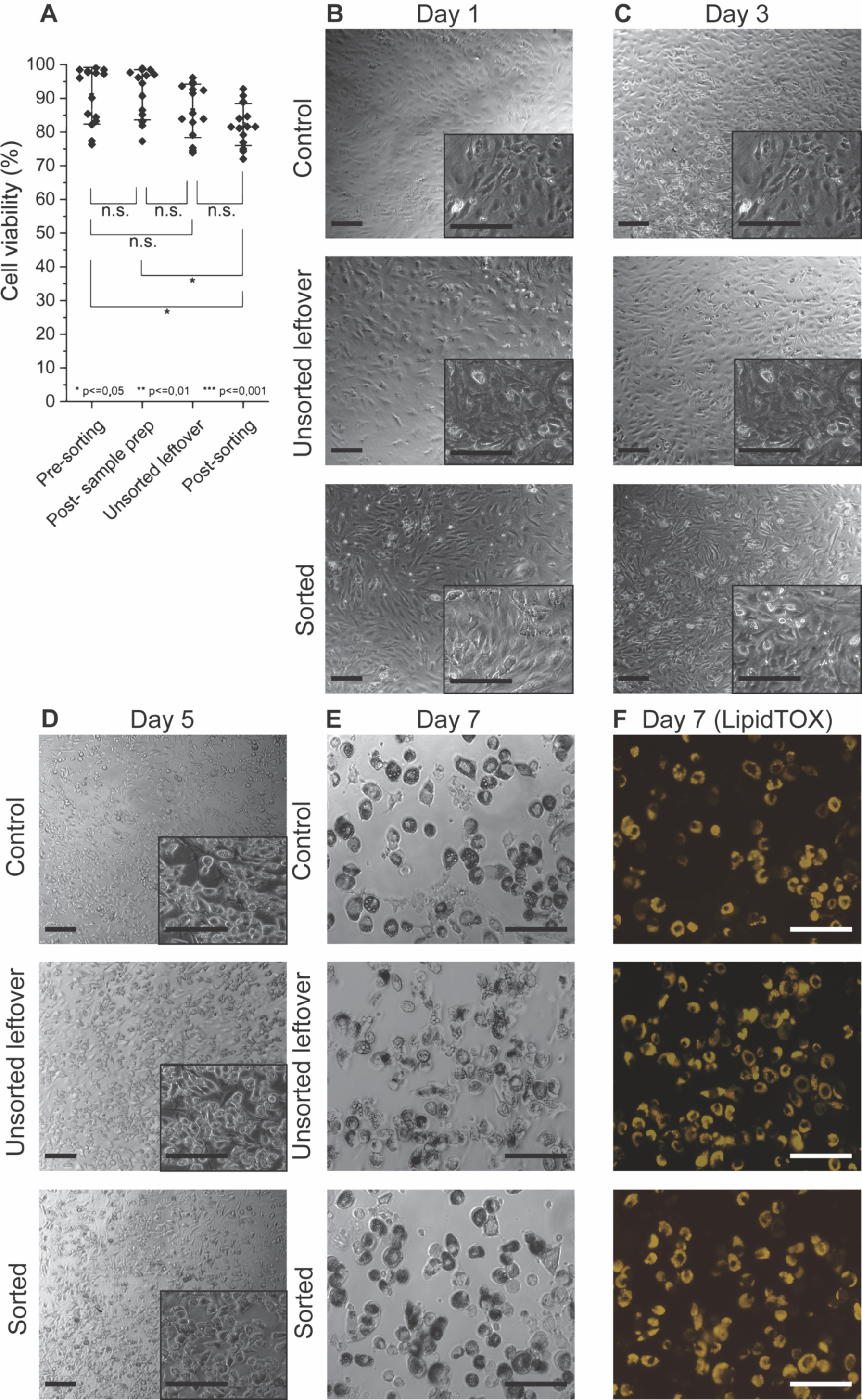
Cell viability and functionality of OP9 cells after MarrowDLD sorting. (A-E) Time-sequential images at different days post-plating following MarrowDLD sorting of undifferentiated OP9 cells. Phase contrast and fluorescence (Yellow, LipidTOX staining) images for control cells (unsorted), unsorted leftover (unsorted sample residual at the inlet reservoir after 3h sorting), and sorted (collected at the «small cells » fraction outlet) after induced adipocytic differentiation for 6 days. Scale bar: 100 µm. (F) Cell Viability measurements: (i) pre-sorting, (ii) post-sample preparation, (iii) unsorted leftover, and (iv) cells collected at outlets post-sorting (small and large cell fractions). Statistical significance was evaluated by the Student’s t-test for independent samples (n=15 MarrowDLD sorting experiments on induced-OP9 samples).

We further assessed post-sorting viability and functionality by comparing sorted progenitor cells (undifferentiated-OP9) to unsorted leftover cells and control cells that were neither sorted nor differentiated (**Fig. 4B-F**). Undifferentiated cultures were chosen for these experiments to ensure adherence for downstream differentiation assays, as otherwise, the low adherence displayed by mature adipocytes from induced-OP9 cultures would have made the sorted cell fractions very difficult to compare. Time-lapse phase contrast microscopy and fluorescence (LipidTOX) images were captured on days 1, 3, 5, and 7. Notably, the sorted OP9 cells, unsorted leftover cells, and unsorted control cell fractions all adhered and proliferated at days 1 and 3, which further confirms post-sorting viability. To determine functionality, we induced differentiation of the three samples. Subsequent imaging on days 5 and 7 revealed clear evidence of adipocytic differentiation, confirmed by LipidTOX staining, which reflected lipid droplet accumulation. Therefore, neither the sorting nor the sample preparation and stirring affected the ability of OP9 progenitors to differentiate in BMAds. Overall, we could thus validate the reliable performance of MarrowDLD in isolating different phenotypes of induced-OP9 adipocytic cells with preserved viability and functionality post-sorting.

### Comparison with FACS sorting

We compared the performance of MarrowDLD with FACS sorting, the gold-standard approach for precise cell sorting at high throughput. FACS sorting relied on LipidTOX intensity to isolate adipocytic cultures into three distinct populations after gating on single viable cells (PI-negative or DAPI-negative), classified respectively as Low, Medium, and High LipidTOX fractions. Note that the FACS-sorting gates for the High LipidTOX fractions were inclusive of all detectable FSC/SSC high events. This gating approach (**Fig. S3A-B**) was implemented employing two different FACS instruments: BD FACSAria™ III (BD Biosciences) at a nozzle pressure of 20 psi and MoFlo Astrios EQ (Beckman Coulter) at a reduced pressure of 10 psi. Lower pressure is expected to preserve fragile adipocytes but implies a slower sorting process.

Following FACS sorting, all output fractions were analyzed by ImageStream to extract the diameter of single viable cells (**Fig. S3C**). **Figure 5** shows the sorting outcomes with Aria and Astrios FACS instruments, with cell size distributions in **Fig. 5A** and **5C**, respectively. Representative ImageStream micrographs of the MarrowDLD fractions are shown in **Fig. 3C**. A single FACS sorting experiment is directly compared with a single MarrowDLD sorting of induced-OP9s with the same passage number, to compare samples with similar size dynamics. The unsorted cells were subjected to the same sample preparation protocols as the sorted fractions and kept at the input reservoirs for the whole process for both MarrowDLD and FACS sortings.

**Figure 5:**
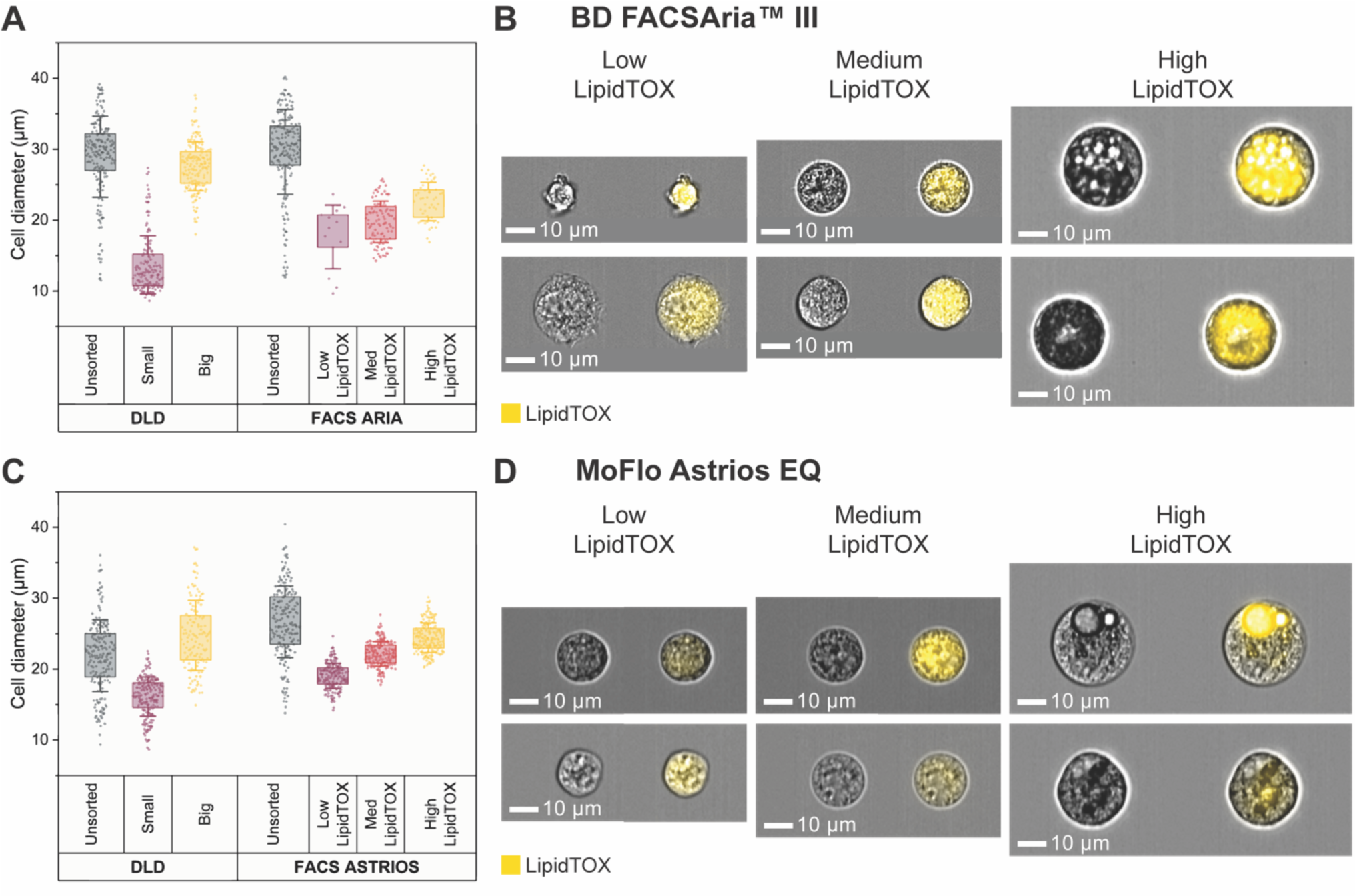
Comparison between MarrowDLD and FACS sorters BD FACSAria™ III and MoFlo Astrios EQ. MarrowDLD size-based separation is compared to FACS sorting based on LipidTOX staining for lipidic content. (A,C) Post MarrowDLD or FACS sorting, cells from the unsorted sample and the sorted fractions were imaged by ImageStream flow cytometry. Distributions of sizes of single viable cells of each fraction after sorting by MarrowDLD, FACS Aria, and MoFlow Astrios FACS instruments. Cell diameters are extracted by ImageStream analysis for each method (200 cells per group or total cells analyzed in the fraction plotted from a single experiment, selected from n=2 biological replicates using BD FACSAria^TM^ III cell sorter and n=3 biological replicates using MoFlo Astrios EQ cell sorter). Data are displayed with standard deviation as error bar, and box plot representing the 25% to 75% data points of the total sample. (B, D) Representative ImageStream images of brightfield and LipidTOX-stained cells from the FACS sorted fractions obtained by FACS Aria and MoFlo Astrios, respectively.

For the FACS Aria versus MarrowDLD comparison, unsorted cells were highly differentiated, with 75% cells above 26 μm (**Fig. 5A-B**). MarrowDLD effectively isolated cells above the predetermined 19 μm cutoff, purifying mature adipocytes and even preserving cells larger than 35 μm. Conversely, the high LipidTOX fraction obtained after FACS Aria sorting surprisingly lacked cells above 30 μm. Notably, we did not observe any intact unilocular adipocytes after sorting by FACS Aria. Despite ImageStream confirming the higher degree of lipid accumulation within the high LipidTOX fraction, as well cells in the other two sorted populations were to a lesser extent positive for the lipophilic stain and overlapped in terms of size. We thus concluded that LipidTOX gating alone was insufficient to discriminate between different degrees of lipidation and that the vast majority of large mature adipocytes was lost during the FACS Aria sorting process.

We then moved to compare the MarrowDLD device to the low-sorting-pressure MoFlo Astrios FACS instrument. We included FSC/SSC gating into the FACS-sorting strategy to better discriminate the Low and Medium LipidTOX populations [14], [47], [49]. For this set of experiments, the unsorted induced-OP9 samples in the MarrowDLD experiment exhibited a narrower range of cell sizes (**Fig. 5C**), therefore including a lower proportion of large, mature adipocytes than for **Fig. 5A**. As shown in **Fig. 5C**, upon ImageStream analysis we found that MarrowDLD isolated cells above the 19 μm separation cutoff and preserved cells larger than 35 μm. For the MoFlow Astrios FACS-sorted fractions, we observed LipidTOX-positive cells in all three fractions, and a harmonious increase in size proportional to the lipid signal (**Fig. 5D**). The MoFlo Astrios sorter could retrieve more intact viable high-LipidTOX cells than the FACS Aria instrument, but again produced losses of large high-LipidTOX cells (**Fig. 5C**). Specifically, cells within the MoFlo Astrios high-LipidTOX fraction displayed lipid droplet accumulation but often did not exhibit complete differentiation, and unilocular adipocytes were not observed (**Fig. 5D**). We, therefore, concluded that contrary to the MarrowDLD device, and although less damaging, the low-pressure MoFlow Astrios FACS sorting still lacked the gentleness required to preserve all fragile adipocytes.

Then, we compared the MarrowDLD performance to the MoFlow Astrios FACS instrument in terms of processing time and yield (**Table 1**). While MarrowDLD is label-free, FACS requires LipidTOX staining incubation (30 minutes) and washing (15 minutes) steps before sorting. Additionally, centrifugation for medium exchange to a FACS buffer potentially harms the cells and introduces buoyancy-based biases. To sort 1.5×10^6^ cells, i.e. the sample size of induced-OP9 cells in a T25 culture flask, MarrowDLD required a 3-hour process, while FACS Astrios approximately 2 hours. Including sample preparation, MarrowDLD took approximately 185 minutes, and FACS required 170 minutes, both within the same order of magnitude.

**Table 1:** Performance comparison. Considering a sample mixture of 1,5×10^6^ induced-OP9 cells (T25 cell culture flask post-differentiation) entirely processed by MarrowDLD or FACS Astrios to isolate mature adipocytes, the table summarizes for each sorting method (i) the sample preparation time, (ii) the total processing time including sample preparation and sorting, (iii) the percentage of large cells lost in the process compared to the unsorted sample calculated as the percentage of unsorted cells above the mean+standard deviation in the high LipidTOX fraction, and (iv) the dilution introduced by the sorting process.

|  | MarrowDLD | FACS |
| --- | --- | --- |
| **Sample preparation time** | 5 minutes (trypsin+ add Optiprep+filter) | 50 minutes (trypsin+ stain LipidTOX/DAPI, wash, filter) |
| **Total processing time** | 3 hours (full sample) | 2 hours (full sample) |
| **Large cell loss** | 7% | 50% |
| **Dilution** | 40x | No dilution (FACS yield 10%) |

To estimate losses of large adipocytes, we compared the size distributions of sorted and unsorted cells (**Fig. 5C**). MoFlow Astrios FACS sorting could not recover 50% of cells larger than 19 μm in the original sample, calculated as the percentage of unsorted cells above the mean+standard deviation in the high LipidTOX fraction. In contrast, for MarrowDLD, the comparison between the large cell fraction and the unsorted population revealed that only 7% of the large cells were lost during sorting. Finally, we should note that FACS does not entail a dilution of the original sample, while MarrowDLD, at this stage of our design, introduces a 40-fold dilution.

Finally, we tested the effectiveness to isolate large lipidated cells from the induced-OP9 single-cell suspensions using the floatation-based protocol developed to isolate primary adipocytes from femoral surgical debris, introduced by *Attané et al.* [19]. Unfortunately, and as reported by the authors (personal communication) we were not able to obtain a floating layer of high-buoyancy adipocytes from our *in vitro* differentiated murine stromal cultures even after long waiting times post-centrifugation (**Supplement. Fig. S4**). The expected cell diameter heterogeneity of induced-OP9 adipogenic cultures was retrieved within the pelleted fraction, including numerous cells larger than 35 μm. Therefore, although we could not compare the buoyancy method for mature adipocyte isolation, we could demonstrate that centrifugation enables the concentration of mature OP9-derived adipocytes to counteract the dilution introduced by MarrowDLD.

### MarrowDLD sorting of spontaneous OP9, induced MSOD, and induced megakaryocytes

MarrowDLD consistently demonstrated high-performance sorting of induced-OP9 adipocytes. To further validate the versatility of our device for fragile cell types, we conducted MarrowDLD sorting experiments on two additional adipogenesis models: spontaneously differentiated OP9 cells, where mature adipocytes are rare (**Fig. 6A**) and adipogenesis-induced MSOD human-derived stromal cells (**Fig. 6B**), as well as human megakaryocytes derived from primary CD34+ cells (**Fig. 6C**). For the adipocyte models, we followed the same sample preparation protocol and sorted an equal number of cells (1.5×10^6^ cells) in each scenario.

**Figure 6:**
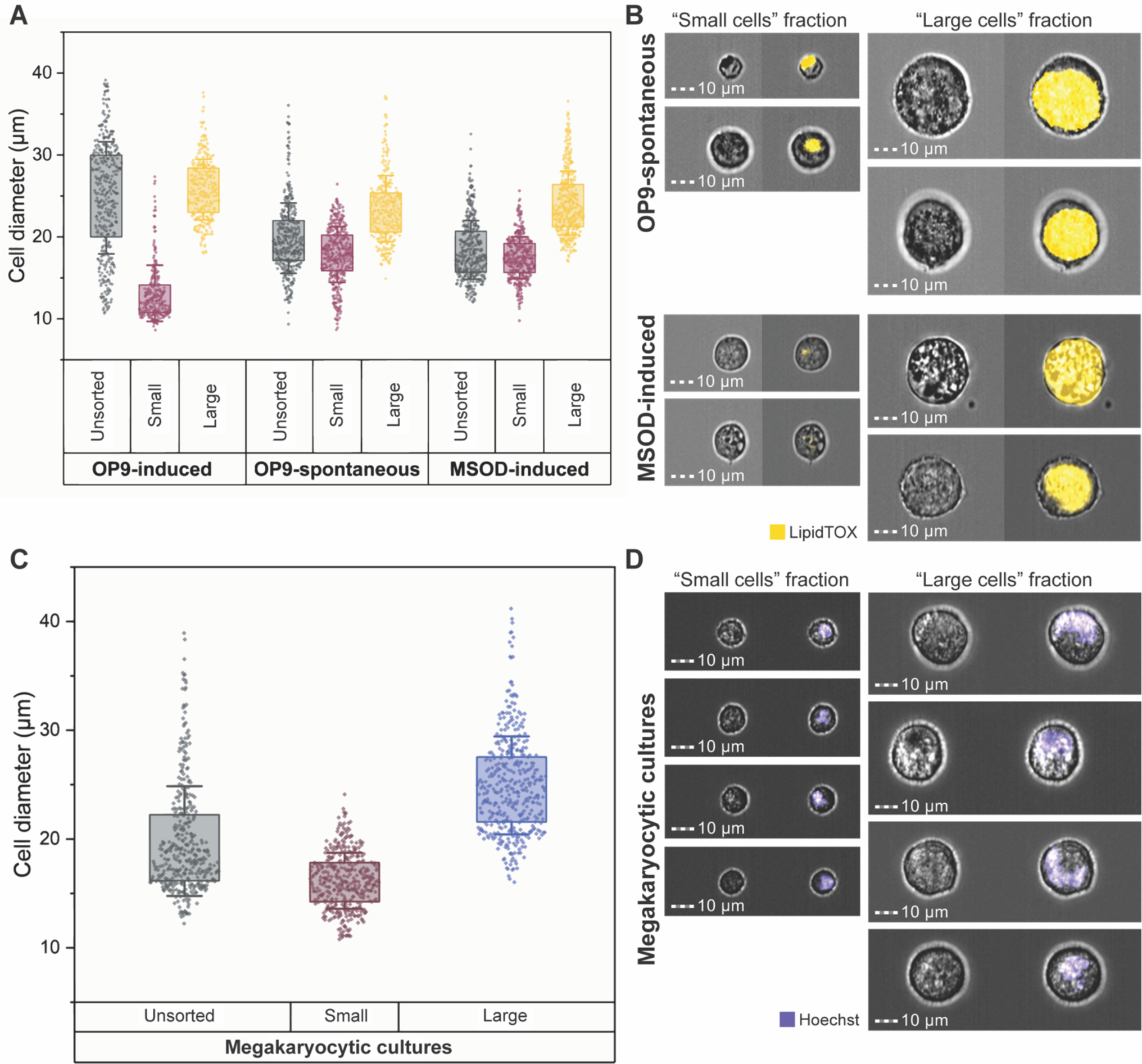
MarrowDLD sorting of different BM cell types. We compared the MarrowDLD sorting of adipocytes from (i) induced (OP9-induced) or (ii) spontaneously differentiated (OP9-spontaneous) mouse-derived OP9 stromal cells, as well as (iii) human-derived MSOD progenitors (MSOD-induced). (A) After MarrowDLD sorting, cells were stained by DAPI and imaged by ImageStream flow cytometry to quantify the diameter of DAPI-negative single cells (n=2 experiments for each cell type, with 400 cells per group). (B) Representative ImageStream micrographs of OP9-spontaneous and MSOD-induced cells (brightfield and LipidTOX-stained) separated by MarrowDLD sorting onto « small cells » (left panels) and « large cells » (right panels) and outlet based on a predefined critical size cutoff of 19 μm. (C) MarrowDLD sorting of human megakaryocytes derived from primary CD34+ hematopoietic progenitor cells. Cells were stained by Hoechst and imaged by ImageStream flow cytometry, to quantify the diameter of the single Hoescht-positive cells (n=2 experiments, with 400 cells per group). (D) Representative ImageStream micrographs upon MarrowDLD sorting of megakaryocytic cultures with a predefined critical size cutoff of 19 μm, showing megakaryocytes on the « large cells » fraction (brightfield and Hoechst-stained after MarrowDLD sorting.

As expected, the post-differentiation single cell-mixture of spontaneous-OP9s, or induced-MSODs, consistently included fewer mature BMAds as compared to the previously tested induced-OP9 samples. However, even if both OP9- and MSOD-derived mature BMAds were rare in these conditions, the MarrowDLD device successfully separated at high purity large cells above the 19 μm separation cutoff (**Fig. 6A**). Specifically, spontaneous-OP9 adipocytes were collected at 90% purity and induced-MSOD adipocytes at 94% purity. ImageStream assessment of the LipidTOX-labeled fractions confirmed that cells within the « small cells » fraction corresponded to the phenotypes of unlipidated progenitor cells or early-stage lipidated cells, while fully-lipidated cells with mature BMAd phenotype were collected within the « large cells » fraction (**Fig. 6B**). This correlation was evident from both the intensity and area of the LipidTOX stain, as well as the cell size inferred from the brightfield images. Thus, we can conclude that MarrowDLD could be also used for high-purity sorting of rare BMAds, from both mouse and human progenitor cells. Notably, the largest adipocytes (> 35 μm) were preserved even if extremely rare.

Finally, we tested our method to isolate differentiated megakaryocytes from primary human progenitors. Differentiation of human CD34+ hematopoietic stem and progenitor cells isolated from peripheral blood was induced *in vitro*. Upon megakaryocytic differentiation, progenitors undergo endomitosis resulting in large, polyploid cells, expressing the characteristic surface markers CD41 and CD42b [10]. The extent of differentiation is largely donor-dependent. Similar to OP9 and MSOD cell lines, the post-differentiation sample contained a mixture of progenitors and various stages of differentiation. As expected, the larger cells as defined by FSC on flow cytometry were double positive for the megakaryocyte-specific markers CD41 and CD42b (**Supplement. Fig. S5**). We tested the capacity of the MarrowDLD device (19 µm separation cutoff) to sort hematopoietic cell mixtures after megakaryocytic differentiation (**Fig. 6C**). Cells above the separation cutoff were isolated at 96% purity and retrieved intact, including cells larger than 35 µm in diameter. Congruently, we found higher polyploidy, as measured by DNA dye Hoechst 33258, in the « large cells » fraction as compared to the « small cells » fraction of the respective MarrowDLD outlets (**Figure 6D**). Overall, our results confirm the robustness and adaptability of MarrowDLD as a powerful sorting method for various fragile cell types, based on a predefined size cutoff, including terminally differentiated cells from human sources.

## Discussion

Our study introduces MarrowDLD (**Fig. 2**), a size-based microfluidic cell sorting device to isolate BM fragile cell types, including adipocytes and megakaryocytes. This method offers high-throughput sorting with low mechanical stress, through a gentle continuous-flow system operated at significantly lower pressures than traditional flow cytometry. Real-time imaging capabilities enable inherent quality control during sorting, valuable for cell biology research, particularly in studying bone marrow adiposity, where assessing minimal BMAds purity remains challenging for standardization and comparability between different studies [2].

To validate the efficiency of MarrowDLD, we first defined the optimal cutoff size to obtain BMAds, and then isolated at high purity OP9-derived BMAds obtained from *in vitro* culture in adipogenic conditions. Despite the induced-OP9 samples comprising a mixture of differentiation stages with varying proportions, MarrowDLD consistently isolated intact cells above the 19 μm separation cutoff within the « large cells » fraction at 97% purity, regardless of the original composition (**Fig.3C**). A separation cutoff of 24 μm was efficient for finer isolation of the largest cells in the mixture.

Microscopy and ImageStream flow cytometric imaging confirmed that all species within the MarrowDLD « large cells » fraction indeed showed lipid accumulation. However, when comparing the two fractions based on LipidTOX signal by ImageStream, we observed that even the small progenitor cells were positively stained, despite the large adipocytes showing a brighter signal. These observations indicate that BMAds sorting methods relying solely on neutral lipid dyes[28]–[30] lack specificity towards mature adipocytes, due to concurrent staining of cell membranes and to the impact of lipid droplets coalescence on fluorescence intensity, as previously suggested by *Hagberg et al.* [14]. While their method demonstrated efficient size-based FACS isolation of large unilocular adipocytes, it required a larger nozzle, reduced pressure, and additional filters, which are often unavailable for standard FACS instruments. Most importantly, despite the adapted shear stress conditions, this process cannot retrieve unfixed adipocytes.

In contrast, our size-based sorting approach offers a non-destructive process preserving in the combined fractions the original cell size distribution in the mixture, enabling the recovery of intact cells for analysis and culture post-sorting. The viability of induced-OP9s fractions after MarrowDLD sorting remained on average consistently above 80% (**Fig. 4A**). Previous studies on DLD devices reported even higher (>95%) DLD post-sorting viability for skeletal stem cells [41] or CTCs [43]. Mature adipocytes are, however, notoriously more fragile. Moreover, we observed a comparable slight reduction of cell viability on leftover unprocessed samples after a few hours of experimental time, indicating that the sorting method itself is purely non-destructive, while adipocyte viability is dependent on the processing time. Mechanical stress from sorting did not affect cell functionality, as progenitor OP9 cells were able to proliferate post-sorting and differentiate into adipocytes (**Fig. 4B-F**). The simple sample preparation, the stirring procedure, and the sample conditions during sorting could be further optimized to improve viability if pertinent. Nonetheless, MarrowDLD as tested already enables the retrieval and replating of very large OP9 BMAds post-sorting, sufficient for further functional studies such as co-culture of BMAds populations with a precisely defined size range, for example, in combination with defined hematopoietic progenitor populations.

We compared our method’s performance to two different FACS instruments, BD FACSAria™ III and MoFlo Astrios EQ (**Fig. 5**). The first operates at 20 psi 100 μm nozzle pressure and the second at 10 psi with the same diameter. Coherently, in our experiments, FACSAria preserved fewer large BMAds than MoFlow Astros. However, even the latter instrument introduced losses as high as 50%. The protocol introduced by *Hagberg et al*. [14] suggested the use of a larger nozzle (150 μm) at 6 psi, however, these settings were not available for our setups, as is expected for most machines in research labs. MarrowDLD, on the other hand, preserved even the large adipocytes and separated the original mixture into two distinct subpopulations with high reproducibility. From a future perspective, we could potentially sort more than two fractions using the same approach with minimal design adaptation. The present MarrowDLD setup comprehends very little instrumentation and easy implementation within a cell and experimental research laboratory. It requires a three-channel pressure pump to manually adjust the inlet pressures and can be readily combined with any microscope used in research labs for cell imaging.

Mature BMAds from tissue biopsies are usually separated by floatation due to their difference in buoyancy [19]. Yet we could not isolate OP9 BMAds by this method. We envision that floatation could be optimized for bulk enrichment of primary BMAds, but not to precisely isolate a pure population of mature BMAds based on size. In our hands, gentle centrifugation enabled to concentrate all differentiation stages in the pellet, which was useful to compensate for the dilution introduced by the MarrowDLD process. At the present stage, our device introduces a 40X dilution that could be reduced by design optimization. Our protocol did not require any buffer exchange as the cells were sorted in their original culture media and could be directly replated post-sorting.

Further to the validation of MarrowDLD sorting of induced-OP9 adipocytes, we tested the device with spontaneously differentiated OP9 samples, where mature adipocytes are rarer and need further enrichment. Even in this case, MarrowDLD consistently isolated the desired fraction with a reproducible separation cutoff achieving 90% purity. Our method was further tested with the MSOD human cell line, where we could retrieve intact MSOD BMAds at 94% purity. Going beyond bone marrow adipocytes, MarrowDLD was effective in sorting megakaryocytes, another notoriously difficult-to-isolate large and fragile cell type in the BM microenvironment. Cells above the separation cutoff were collected at 96% purity within the large cell fraction and showed characteristic MK polyploidy.

Considering a sample of 1.5 million cells, MarrowDLD showed a comparable processing time to FACS (**Table 1**). The ease of sample preparation without labelling, washing, or buffer exchange, contributed to a substantial reduction in the total processing time. Moreover, we operated the microfluidics at a relatively large flow rate (up to 1 mL/hour) without compromising cell viability, and we believe that the flow speed could be even increased with acceptable post-sorting functionality. The design of MarrowDLD could also be adapted to have multiple chambers in parallel as an easy strategy to increase the throughput.

Overall, the current MarrowDLD system enables the recovery of at least 10^5^-10^6^ intact and functional fragile BM cells per sorting run, suitable for further studies and post-sorting culture. As a future perspective, we envision that the system could be considered for scale-up towards clinical applications requiring large throughput, and could be adapted to sort multiple differentiation stages from the original mixture. The isolation of pure populations of mature BM cells (BMAds and MKs) with a precisely defined size range offers biologists new possibilities to study how each subpopulation interacts with neighboring cells regulating blood production and blood clotting. Studying the roles of these cells within the bone marrow niche can provide insights and reveal novel therapeutic targets for various hematological disorders, including leukemia, myeloproliferative neoplasms, and thrombocytopenia, as well as diseases affecting bone health.

## Methods

### Mouse-derived mesenchymal stem cell (OP9) culture and differentiation

OP9 cells (provided by T. Nakano, Kyoto University, Japan) were cultured and differentiated as described in Campos *et al*. [49]. Cells plated at 20,000 cells/cm^2^ were maintained in MEM-α with GlutaMaxTM (Gibco, Cat. No. 32561) supplemented with 10% FBS (Gibco, Cat. No. 10101), and 1% penicillin/streptomycin (Gibco, Cat. No. 15140) at 5% CO2 and 37°C. Cells were split using a trypsin-EDTA solution for 5min when subconfluent (80%). A differentiation cocktail composed of culture medium along with dexamethasone (Sigma, Cat. No. D2915, 1*μ*M in ethanol), isobutyl-methylxanthine (Sigma, Cat. No. I7018, 0.5mM in DMSO), and insulin (Sigma, Cat. No. I0516, 5 *μ*g/mL) was used to perform the adipocytic differentiation for 6 days. For spontaneous adipocytic differentiation, OP9 cells were kept in culture for 18 days. Every 3-4 days, half of the medium was replaced with fresh culture medium.

### Human marrow stromal cell (MSOD) culture and differentiation

MSOD cells, an immortalized human bone marrow stromal line, were obtained from Ivan Martin’s laboratory in Basel, Switzerland [50]. MSOD cells were maintained in MEM-α with HEPES (Thermofisher, Cat. No. 15630-056, 10mM). When subconfluent, cells were split as for OP9 cells. A differentiation cocktail, composed of culture medium along with dexamethasone, isobutyl-methylxanthine, insulin, and rosiglitazone (AdipoGen, Cat. No. CR1-3571, 15μM), was used for adipocytic differentiation for 21 days. When cells were seeded, the differentiation cocktail was added at double concentration. On days 6 and 18, half of the medium was replaced with fresh culture medium. On day 12, half of the medium was removed and replaced with a differentiation cocktail.

### Mouse and Human-derived mesenchymal cell staining

#### For flow cytometry sorting and imaging using ImageStream (Cytek Bioscience)

Prior to flow cytometric sorting, cells were stained with LipidTOX^TM^ Deep Red neutral lipid stain (Invitrogen, Ref. No. H34477, supplied as 1000X for standard assays) for 30 min at 37°C and DAPI (Axonlab, Ref. No. A4099.005, 5 mg/mL) for 10 min at room temperature. The cells were subsequently detached by incubating for 5 min in 0.05% trypsin (Gibco, Cat. No. 25300054) at 37°C. The OP9 cells were gently resuspended in PBS before being filtered with a 100μm cell strainer in FACS tubes.

After DLD sorting, the different sorted fractions were stained for LipidTOX^TM^ Deep Red neutral lipid stain (Invitrogen, Ref. No. H34477, supplied as 1000X for standard assays) for 30 min at 37°C and DAPI (Axonlab, Ref. No. A4099.005, 5 mg/mL) for 10 min at room temperature prior subjected to Imagestream.

#### For fluorescence microscopy imaging

At different time points during the differentiation time course, plated cells were stained with live fluorescence dyes: BODIPY™ (boron-dipyrromethene, Invitrogen Ref. No. D3922,10ng/mL), or LipidTOX^TM^ Deep Red neutral lipid stain (Invitrogen, Ref. No. H34477, supplied as 1000X for standard assays) for 30 min at 37°C. Cells were incubated with the dyes in FluoroBrite phenol red-free DMEM medium (Gibco, Cat. No. A1896701) supplemented with 10% FBS and 1% penicillin-streptomycin for 30 min at 37 C in the dark, washed twice with warm PBS and imaged in FluoroBrite medium using EVOS 5000 imaging system (ThermoFisher, Cat. No. AMF5000).

### Human-derived megakaryocyte differentiation from primary progenitors

All experiments were performed in accordance with relevant named guidelines and regulations, and in accordance with the Declaration of Helsinki. Human blood cells were obtained from discarded blood cell obtained from anonymous healthy blood donors at the Red Cross Inter-regional Transfusion Center (Epalinges, Switzerland), and their use approved by the competent authority on research involving humans (Cantonal Ethical Committee at Vaud canton, CER-VD, Switzerland). Informed consent was obtained by the Inter-regional Transfusion Center from all participants before collection. Human CD34+ cells were isolated from discarded buffy coats obtained upon blood unit preparation by magnetic-activated cell sorting (MACS). Briefly, the buffy coat collected at the blood transfusion center was diluted in an equivalent volume of PBS 2mM EDTA. 15mL Ficoll Plaque (GE, Cat. No. 17-1440-03) was added in a 50mL tube. 30mL of the blood mixture was then added delicately on top of Ficoll. Cells were then centrifuged for 30 minutes at 400g without brake and washed with PBS 2mM EDTA. Cells were treated with red blood cell lysis buffer (BioLegend) for 2 minutes and washed with PBS. For CD34+ cell isolation, cells were subjected to staining according to manufacturer protocol (CD34 microbead kit: Microbeads Miltenyi, Cat. No. 130-100-453).

Isolated CD34+ cells were seeded at a density of 50,000 cells/mL in Expansion Medium for 7 days. Expansion medium was composed of StemSpan Serum Free Expansion Medium (Stemcell Technologies, Cat. No. 09650) supplemented with 20ng/mL StemSpan Megakaryocyte Expansion Supplement (Stemcell Technologies, Cat. No. 02696), 20ng/mL human plasma-derived low-density lipoprotein (hLDL) (Stemcell Technologies, Cat. No. 02698), 1uM StemReginin1 (SR1) (Biogems, Cat. No. 122499) and 1% GibcoTM Penicillin-Streptomycin-Glutamine (PSG) (Thermofisher, Cat. No. 10378016). At day 7, cells were centrifuged and seeded at a density of 100,000 cells/mL in a differentiation cocktail for 7 days. The differentiation cocktail was composed of StemSpan Serum Free Expansion Medium supplemented with 20ng/mL hLDL, 1uM SR1, 0.5ug/mL human recombinant Thrombopoietin (TPO) (StemCell Technologies, Cat. No. 78210), and 1% PSG (Thermofisher, Cat. No. 10378016). Successful Differentiation was confirmed by flow cytometry using CD41 (APC, clone HIP8, Biolegend Cat. No. 303710, Dilution 1:200) and CD42b (PE, clone HIP1, Biolegend, Cat. No. 303906, Dilution 1:200) surface detection. Briefly, 20uL of the cells were diluted with up to 100uL with PBS. Cells were then stained with CD41 and CD42b for 20 minutes and subjected to flow cytometry (Accuri C6 PLUS, BD).

After DLD sorting, the different sorted fractions were stained for polyploidy (Hoechst 33258, Invitrogen H3570). Briefly, cells were washed in PBS and centrifuged for 5 minutes at 200g. Cells were resuspended in 100uL PBS with 1ug/mL Hoechst 33258. Cells were incubated for 15 minutes at room temperature. Cells were then washed as previously and resuspended in 50uL of PBS prior subjected to Imagestream (Cytek Bioscience).

### Sample preparation for MarrowDLD

The device operation was first assessed by sorting polystyrene beads of 15, 18, and 30 µm in diameter (Spherotec Inc.). Beads are suspended in 1xPBS supplemented with 1% (w/v) BSA (Sigma, Cat. No. A7906) to prevent aggregation and adhesion to the channel walls.

Both OP9 and MSOD BMAds, as well as megakaryocytes, were sorted by MarrowDLD following the same sample preparation protocol. Cells were washed, detached, and resuspended in culture media supplemented with 16% of Optiprep (StemCell, Cat. No. 07820) to ensure a single-cell suspension and filtered with a 100μm cell stainer (PluriSelect, Cat. No. 43-50150-01) for clusters removal. The optimal cell concentration for MarrowDLD sorting was 10^6^ cells/mL.

### Device design and fabrication

Our MarrowDLD sorting module is based on the design proposed originally by *Huang et al.* [35]. The microfluidic device has a single central inlet for sample delivery and two side ones for cell stream focusing. Two outlets collect two separated fractions. Microfluidic resistors were placed at the inlets to calibrate the applied pressure and at outlets for even splitting of the output flow. The DLD array was designed to provide a specific critical size for sorting according to the empirical model by *Davis et al.* [39]. We designed and fabricated devices at nominal critical sizes: 15, 17, 20, 22.5, and 25 μm. The nominal critical sizes of 15 and 20 resulted in a separation cutoff of respectively 19 μm and 24 μm for induced-OP9s sorting.

The microfluidic device consisted of a polydimethylsiloxane (PDMS) module bonded to a glass substrate. The PDMS casting mold was fabricated by photolithography patterning of 50 μm SU-8 resist (Microchem 3025, Microresist Technologies, Berlin, Germany) on silicon. Surface conditioning of the silicon/SU-8 mold by silanization with Chlorotrimethylsilane (TMCS, Sigma Cat. No. 386529) aided the PDMS release. After silanization, a mixture of PDMS precursor and crosslinker (SYLGARD™ 184 Silicone Elastomer Kit, Dow Corning Corp.) at a 1:9 ratio is dispensed onto the mold and cured by heating at 80°C for 2 hours. Finally, the PDMS module is activated by oxygen plasma (550 mTorr, 29 W, for 45 s) and bonded onto a glass coverslip. The bonding is accelerated by a 2-minute baking step at 80°C.

### MarrowDLD sorting

Vials were connected to inlets and outlets by Tygon® tubing (Cole-Parmer, Cat. No. GZ-06420-02) paired with metallic connectors (Unimed, Cat. 200.010-A). Inlet reservoirs were paired with a pressure pump (Fluigent, Flow EZ, Cat. No. LuFEZ-1000) driving the flow. The chip was primed with cell culture media overnight before operation. After sample preparation, the mixture was delivered to the central inlet reservoir and kept under magnetic stirring to ensure homogeneity. To avoid clogging issues throughout the whole sorting experiment (1h to 5h), a continuous filtering method was developed by embedding a 100 μm cell strainer membrane at the tube extremity. The device is mounted on a microscope (Leica DM IL LED) equipped with a camera (ORCA-Flash4.0 V3 Digital CMOS camera) to visualize the cells trajectories in real-time. After sorting, the two fractions were collected for further analysis.

### Statistical Analysis

Results are displayed as Mean ± SD, unless otherwise stated, by Origin 2022 (OriginLab Corporation, Northampton, MA, USA). Two-tailed Student’s t-test for independent samples was used to test statistical significance, after Shapiro–Wilk normality test.

### Data availability statement

The datasets generated and analysed during the current study are available in Zenodo repository, https://doi.org/10.5281/zenodo.8318213

## Acknowledgements

The authors acknowledge financial support from the Swiss National Science Foundation (205321_179086) and (PP00P3_183725), the ETH Personalized Health and Related Technologies Initiative (PHRT-308), Innosuisse (56547.1) and the University of Lausanne (UNIL). We thank the Center of MicroNanoTechnology (CMi, EPFL) for assisting the microfabrication process, the BioImaging and Optics Platform (BIOP, EPFL) for the access to microscopy instruments, and the Flow Cytometry Facility (UNIL), in particular Danny Labes and Francisco Sala de Oyanguren for the FACS sorting and ImageStream analysis, respectively. We express gratitude to Marjolaine Boulingre for her help in the early stages of the project.

## Author contributions

G.P., R.S., C.G., and O.N. conceived the concepts of the research and designed the experiments. G.P. designed the MarrowDLD system architecture. G.P., C.O., L.G., and F.A. microfabricated the devices. G.P., R.S., C.O., L.G., and F.A. performed the experiments including MarrowDLD sorting, Imagestream analysis, FACS sorting, cell culture and functionality assays. G.P. analyzed the data. M.H. collected human blood samples and derived MKs from in-vitro differentiation. G.P. and R.S. co-wrote the manuscript; all the authors revised the manuscript.

## Additional Information

The authors declare no competing interests. Correspondence and requests for materials should be addressed to G.P. or R.S.

## Notes

https://doi.org/10.5281/zenodo.8318213

### Competing Interest Statement

The authors have declared no competing interest.

