## Supplemental Information for "MarrowDLD: a microfluidic method for label-free retrieval of fragile bone marrow cells"

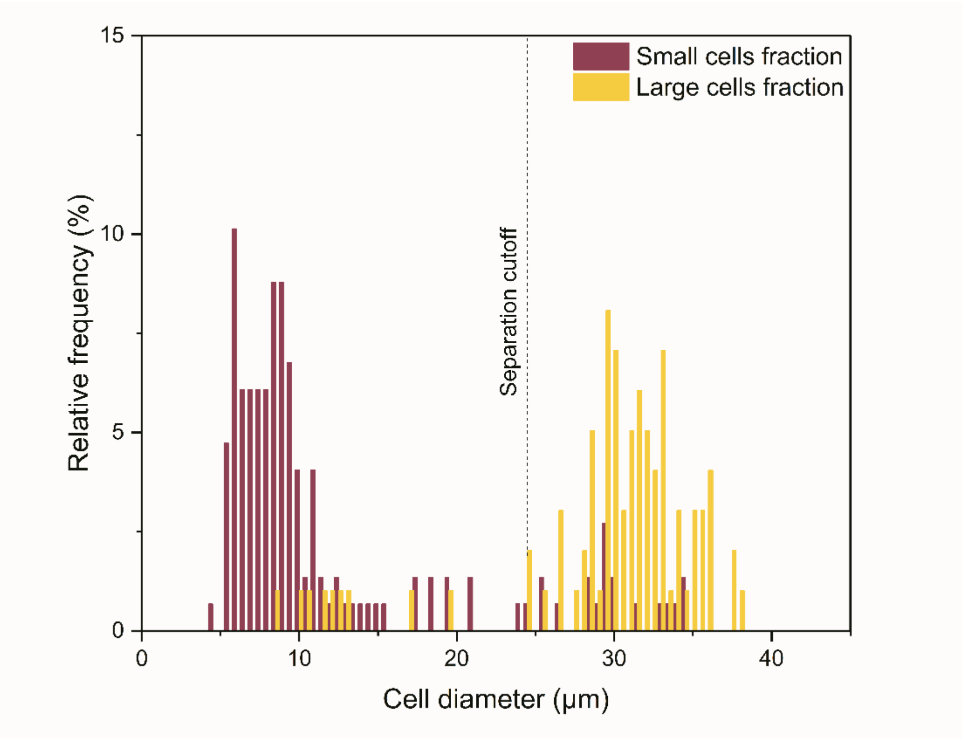


**Figure S1: Sorting of induced OP9 cells with a separation cutoff of 24 μm.** Quantification of cell diameter distributions in each outlet ("small cells" fraction, violet; "large cells" fraction, yellow) after sorting of induced-OP9 cells by means of the MarrowDLD system with a 24 μm separation cutoff (24 μm critical size). The quantification was based on phase contrast microscopy measurements of cells collected at the outlets and replated for imaging after sorting. Cell diameter measurements were performed using QuPath0.3.2 software. The cell diameter distributions were displayed as relative frequencies over 150 cells per sample group from n=2 sorting experiments. The same protocol for purity quantification and empirical characterization of the critical size was performed for MarrowDLD devices with a separation cutoff of 19 μm as displayed in Figure 3C in the main text.

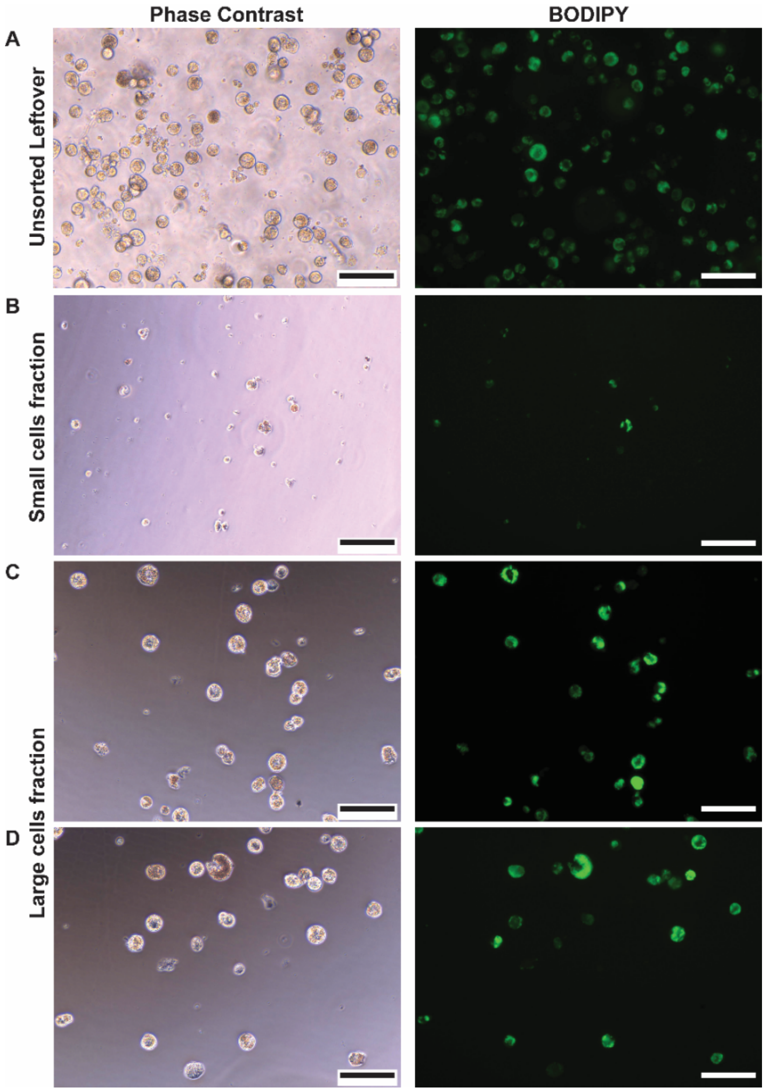


**Figure S2: Induced OP9 cell phenotype visualization post-DLD cell sorting.** Phase contrast (left) and fluorescent (BODIPY, right) images of induced OP9 post-sorting plating. (A-D) Cells at day 6 post-differentiation and DLD post-sorting (stirred unsorted remaining in the inlet [A], cells from the small cells fraction [B], and the large cells fraction [C-D]) were plated and stained with BODIPY for lipid content visualization (scale = 100µm).

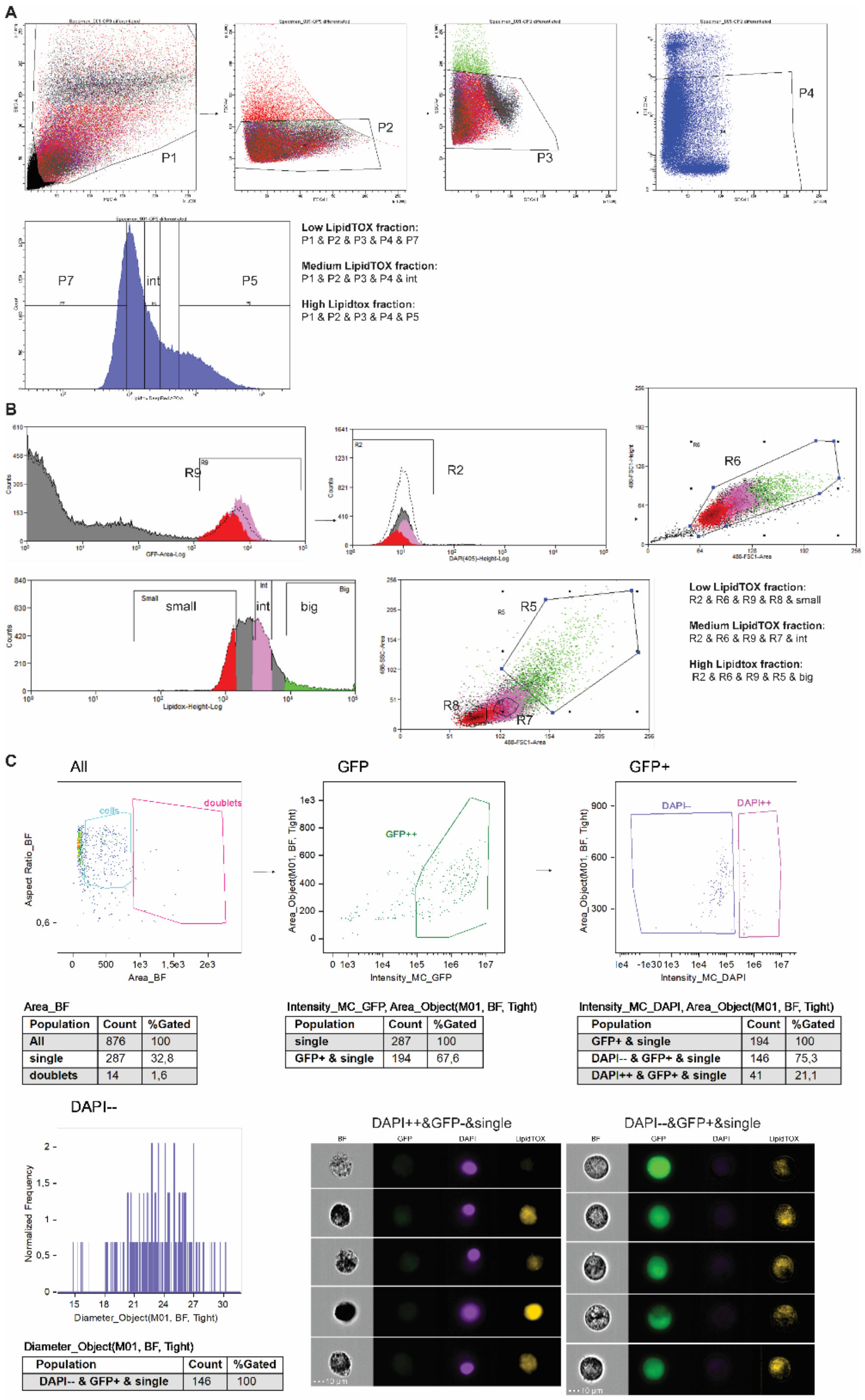


**Figure S3: FACS gating strategies for LipidTOX-based sorting of a sample of induced OP9 cells and gating approach for ImageStream flow cytometry imaging. (A)** FACS sorting using BD FACSAria Cell Sorter. Three subsequent gating steps are performed to isolate singles namely plotting FSC-A vs. SSC-A (P1), followed by FSC-H vs. FSC-W (P2), and finally SSC-H vs. SSC-W (P3). Subsequently, we performed gating for Propidium iodide PI-negative cells (P4), to exclude non-viable cells. In the next step, LipidTOX fluorescence of the viable singlets is evaluated on a histogram: three non-contiguous subpopulations were then sorted based on LipidTOX intensity (here named ‘P7’, ‘int’, and ‘P5’). These three fractions were collected after FACS sorting and analyzed by Imagestream, as displayed in Figure 5.A in the main text. **(B)** FACS sorting using MoFlo Astrios Cell Sorter. In the first step, the GFP-positive cells are selected (R9). The OP9-GFP line, which constitutively expresses GFP in the cytoplasm (REF), was used for these experiments to facilitate the exclusion of debris and non-viable cells through

GFP+DAPI- gating (R2). We then performed singlet gating to exclude cell aggregates based on FSC-A vs. FSC-H (R6). Subsequently, LipidTOX fluorescence of the viable singlets was evaluated on a histogram: three subpopulations are identified based on LipidTOX intensity (here named 'low, 'int', and 'high'). In a final gating step, we restricted the sorted Lipidox low population to the lower FSC/SSC region (R8). To further refine the populations to the most representative stages of adipocytic differentiation, the Lipidox high population was further restricted to the highest FSC/SSC region (R5), and the Lipidox intermediate population was selected to be in-between the other two based on their FSC/SSC profile. The obtained three fractions (namely "Low-LipidTOX", "Medium LipidTOX", and "High LipidTOX" factions) have been collected after MoFlo Astrios FACS sorting and analyzed by Imagestream, as displayed in Figure 5.C in the main text. **(C)** ImageStream gating approach to select single viable cells from FACS-sorted or MarrowDLD-sorted samples. Here a "Large cells" fraction collected after MarrowDLD sorting of OP9-induced samples is shown to illustrate the strategy. First, singlets were selected by plotting the brightfield (BF) area vs. the BF aspect ratio, and GFP-positive cells were selected to better exclude debris and ruptured cells. Subsequently, DAPI intensity vs. BF area is plotted to select viable (DAPI-negative) cells. With this approach, we identify single intact viable cells, of which we obtain the diameter size distribution. This method is applied to all the different populations characterized by Imagestream analysis for extracting the diameter size distribution, namely the data shown in Figures 5A and 5C, as well as Figures 5A and 5C in the main text.

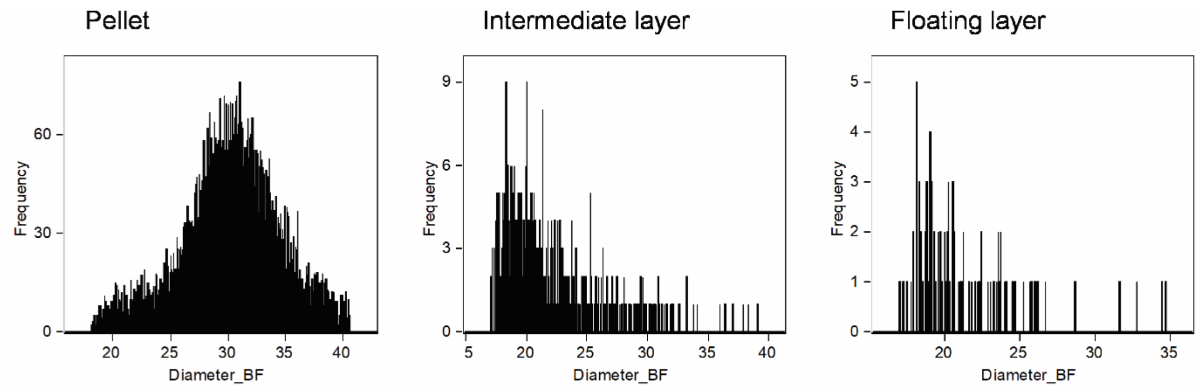


Diameter_BF

| Population | Count | %Gated |
|---|---|---|
| DAPI- & cells | 11106 | 100 |

Diameter_BF

| Population | Count | %Gated |
|---|---|---|
| DAPI- & cells | 485 | 100 |

Diameter_BF

| Population | Count | %Gated |
|---|---|---|
| DAPI- & cells | 107 | 100 |

**Figure S4: Separation by floatation of Induced-OP9.** A sample of induced-OP9 cells was subjected to the floatation protocol described by *Attane' et. al.* Three fractions are collected, namely the bottom layer (pellet), the middle supernatant layer (intermediate layer), and the top supernatant (floating layer), subsequently analyzed by ImageStream flow cytometry imaging. Here we display frequency distributions of cell diameters for the three fractions, as well as the amount of viable (DAPI-) cells collected at each fraction. Cells in the intermediate and floating layers are in negligible number compared to the pellet, indicating that mature BMAds are not separated by floatation and all the different cell types in the mixture are collected in the pellet after centrifugation.

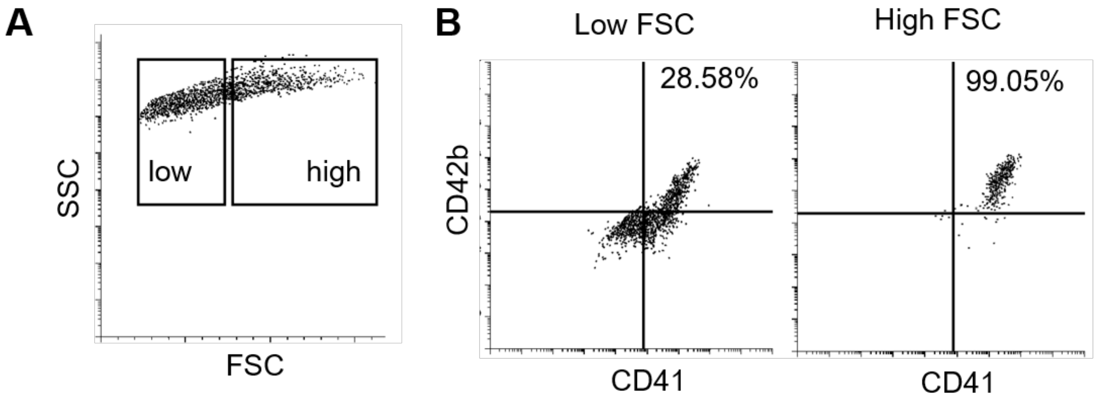


**Figure S5: CD41+CD42b+ megakaryocyte lienage cells are enriched in the "large cells" fraction in CD34+ progenitor cultures.** CD34+ cells isolated from peripheral blood were differentiated towards megakaryocytes for 14 days. Total cells were stained for CD41 and CD42b surface markers and subjected to flow cytometry analysis. (A) Cells were gated according to the FSC low (Smaller fraction) and FSC high (Larger fraction). (B) The larger fraction is enriched in CD41+CD42b+ megakaryocyte (right panel) compared with the smaller fraction (left panel).
